# Integrated high-resolution phenotyping, chlorophyll fluorescence induction kinetics and photosystem II dynamics under water stress and heat in Wheat (*Triticum aestivum*)

**DOI:** 10.1101/510701

**Authors:** Arun K. Shanker, Robert Coe, Xavier Sirault

## Abstract

An experiment was conducted in controlled conditions in three varieties of wheat under water stress, heat and heat +water stress treatments with the objective of studying Chlorophyll a fluorescence, chlorophyll fluorescence induction kinetics and the function of Photosystem II by plant phenotyping as affected by stress. We hypothesised that during stress, specific adaptive strategies are employed by plants, such as structural and functional changes in PS II by which they acquire new homeostasis which may be protective adaptations. Water stress stress treatment was imposed on Water stress and Heat +Water stress treatments at 43 DAS. Heat treatment was imposed on 48 DAS. Maximum quantum yield of primary photochemistry was measured with PAM 2500 and OJIP was measured with FluorPen FP 100 after the onset of stress at four observation times on two days viz., pre-dawn and afternoon during stress. In addition continuous monitoring of photosynthetic efficiency was done with Monitoring PAM. Heat +Water stress stress was more detrimental as compared to Heat or Water stress alone in terms of maximum quantum yield of photochemistry. This could have been due to higher decrease in connectivity between PSII and its antennae resulting in lower photosynthetic efficiency resulting in the impairment and disruption of the electron transport. K step was observed in heat stress and heat +Water stress stress which may be because of damage to Oxygen Evolving Complex indicating that low thermostability of the complex. The stress treatments had a reduction in the plastoquinone pool size as indicated by the reduced area above the OJIP curve. Our study indicated that the instrument PAM 2500 sensed both stresses separately and combined earlier than the other instruments, so in terms of sensitivity PAM 2500 was more effective than FluorPen FP 100 and MultispeQ. Rapid screening of stress was more effectively with FluorPen FP 100 and MultispeQ than by PAM 2500.

## Introduction

Photosystem II (PSII) photochemistry, light energy absorption, excitation energy trapping, and conversion of excitation energy into electron flow can be targets of stress like water stress and heat and reflect on the overall growth and development of the plant (Poudyal 2019, Sasi et al 2018). Plants also respond to stress stimuli and if the stress is not eliminated in time, a rapid damage of the photosynthetic system occurs. To understand the mechanism action of abiotic stress phenotyping individual quantitative physiological parameters – Photochemisty, growth and development, biomass and yield as determined plant phenotyping can help us unravel more complex stress tolerant traits. Chlorophyll fluorescence (ChlF) measurements is now known as an important tool to study stress in plants (Kalaji et al 2016; Hazrati et al 2016; Mathobo et al 2017; Dąbrowski et al 2016)

We hypothesise that specific strategies are employed by plants, such as structural and functional changes in the photosynthetic systems by which they acquire new homeostasis which may aid in protective adaptations. Changes may be as a response to stimuli or alterations to counter stress. Heat and water deficit stress when simultaneously imposed can have effects that may not be seen when individually imposed. Insights into the mechanism of water stress and heat stress can be an important out put of the study and this can help in elucidating the processes that are responses to stress and also the processes that are countering stress in plants (Ergo et al 2018, Zandalinas et al 2018). The objectives of the study was to study the Chlorophyll a fluorescence transient in wheat (*Triticum aestivum*) under water deficit and heat stress, the structure and function of Photosystem II, the details of electron transport chain and its role in acclimation to water deficit and heat stress in wheat (*Triticum aestivum*) and the individual quantitative physiological parameters by plant phenotyping that form the basis for more complex abiotic stress tolerant traits. Specific study objectives was to elucidate the mechanism of multiple abiotic stress in crops with a focus to acquire skills and knowledge to work in High Resolution Phenotyping. The long term outcome of this study will be that it will pave the way for development of climate ready cultivars in wheat and also in other dryland crops.

## Materials and Methods

Wheat seeds of three varieties viz., Federation, Mace and Bolac was soaked with water in a petri dish and kept in dark at 4° C for two days and after germination they transplanted to trays at 7 DAS and then transplanted to pots at 15 DAS. Wheat plant were grown in a controlled environment growth chamber under a photoperiod of 12 hours Day and 12 hours night in pots of 20X20 cm diameter with a capacity of 6.5 L with wheat special soil mix sourced from CSIRO. The photon flux density in the chamber was 300 μ mol. m^−2^.s^−2^ for the first two hours of the day, 400 μ mol.m^−2^.s^−2^ for the next two hours, 500 μ mol.m^−2^.s^−2^ for the next four hours, 400 μ mol.m^−2^.s^−2^ for the next two hours and 300 μ mol. m^−2^.s^−2^ for the last two hours of day to simulate near natural conditions. The temperature in the chamber was 18° C in the first hour and was progressively increased by 1 degree till the 6 hour of day to reach 23° C, then the temperature was reduced progressively by 1 degree to reach 18° C in the last of the day. Major and micro nutrient mixture were applied at appropriate time to avoid deficiency in any Osmacote (Scotts, Australia) of the nutrients. The pots were irrigated every with tap water. Water stress treatment was imposed on water stress and Water stress + Heat treatments at 43 DAS by withholding irrigation. Heat treatment was imposed so as to coincide with water stress treated plants. Heat treatment was imposed on 48 DAS plant in heat and heat + water stress treatments by increasing the temperature to 25 ° C in the first hour of day and 30 ° C and 35 ° C in the second and third hour of day, then the temperature was increased to 37 ° C for four hours after which the temperature was reduced to 35 ° C for the next two hours and 30 ° C for the next hour and 25 ° C for the last hour. The same night temperature of 17 ° C was maintained in all the treatments. Soil moisture content was calculated by the gravimetric method and Table 1 shows the Soil moisture content in the control and treatment pots, water stress and water stress + heat treatments had less than 50 percent of soil moisture of that of the control pots at after 5 days after imposition of water stress stress and 1 day after imposition of heat stress at 49 DAS.

**Table 1.** Soil Moisture Percentage in pot of different treatments in different varieties of wheat at after 5 days after imposition of water stress stress and 1 day after imposition of heat stress at 49 DAS.

| Treatment | Soil Moisture Percentage |
| --- | --- |
| Control | 36.38 |
| Water stress | 18.09 |
| Heat | 35.19 |
| Heat + Water stress | 16.12 |

Relative Water Content was measured according to Smart et al (1974), fully expanded leaves were excised and fresh weight (FW) was immediately recorded from control and stressed plants; then leaves were soaked for 4 h in distilled water at room temperature under a constant light, and the turgid weight (TW) was recorded. After drying for 24 h at 80 °C total dry weight (DW) was recorded. A minimum of one leaf in a treatment mid way after imposition of stress, this one way to quantify stress.

Maximum quantum yield of primary photochemistry was measured with three instruments PAM 2500, FluorPen FP100 and MultispeQ

### Slow Fluorescence kinetics

Dark Adapted leaf - Fv/Fm at two types of Saturation Pulse data which is Fo - Basic chlorophyll fluorescence yield recorded with low Measuring Light intensities. Fm Maximal chlorophyll fluorescence yield when photosystem II reaction centres are closed by a Saturation Pulse Light exposed leaf - Fo’ - Minimum chlorophyll fluorescence yield in the state of open photosystem II reaction centres – this is measured with in the presence of far-red illumination with Actinic Light switched off Fm’ - Maximal chlorophyll fluorescence yield when photosystem II reaction centres are closed by a strong light pulse Ft - The Ft denotes the continuously recorded fluorescence. The value of Ft measured shortly before a Saturation Pulse with light-exposed samples Calculated parameters PS II yield – Fv/Fm and Y (II) ETR - from Y(II) and PAR NPQ - quantification of non-photochemical quenching alternative to qN calculations

### Fast Fluorescence kinetics

The trigger Pattern Poly300ms.FTM will used to measure the OJIP curve The trigger results in a Fast Kinetics which establishes a baseline in the absence of Measuring Light at negative time values. Shortly before time “zero”, high frequency Measuring Light is switched on the establish the Fo level fluorescence. At time zero, a polyphasic fluorescence kinetics is initiated by a strong multi turn-over flash and, simultaneously, the time resolution of measure-ment is further increased by switching to maximum frequency of Measuring Light. Statistical Analysis

The data was analyzed statistically using a general linear model for analysis of variance (repeated measures Anova) (Wilkinson et al., 1996). Significance between control and treatments were compared at 0.05 and 0.01 probability levels

## Results

Relative Water Content (RWC) at 49 DAS in the leaves of wheat varieties Federation, Mace and Bolac in control and stress treatments is shown in Table 2. All the stress treatments water stress, heat and combination stress treatment heat + water stress had reduced RWC content compared to control in all the varieties. The varieties did not exhibit significant differences among themselves under control conditions, with Bolac showing highest RWC of 80.57 followed by Federation and Mace. In the case of water stress treatment the RWC was significantly less compared to control and the variety Federation has significantly less RWC as compared to Bolac which had a highest RWC of 70.19 under water stress. There was no significant difference between Federation and Mace in RWC under water stress. The heat treatment reduced the RWC in wheat as compared to control by a maximum of 6.22 percent in Bolac and in Federation and Mace the reduction was less than 5 percent. The RWC varied significantly under heat treatment as compared to control in the heat treatment in all the varieties, although there was no significant difference between varieties. Under water stress + Heat treatment the lowest RWC was observed in Federation (56.28) which was 23.03 percent less than control. The least difference between control and treatment was observed in Bolac (17.99). RWC in the variety Federation exhibited significant difference from Bolac.

**Table 2.** Relative Water Content in leaves of wheat varieties after 5 days after imposition of water stress stress and 1 day after imposition of heat stress at 49 DAS. The figures in parenthesis indicate SEM, * represents significance at *p*=<0.05

| Variety | Treatment |  |  |  |
| --- | --- | --- | --- | --- |
|  | Control | Water stress | Heat | Heat + Water stress |
| Federation | 79.31 ($\pm 1.40$) | 64.06 ($\pm 0.82$) * | 75.08 ($\pm 1.41$) | 56.28 ($\pm 0.36$) * |
| Mace | 78.75 ($\pm 1.62$) | 66.75 ($\pm 0.64$) * | 73.02 ($\pm 1.29$) | 58.35 ($\pm 1.48$) * |
| Bolac | 80.57 ($\pm 0.78$) | 70.19 ($\pm 0.83$) * | 74.35 ($\pm 0.84$) | 62.58 ($\pm 1.47$) * |

Fig.1 shows the maximum quantum yield of primary photochemistry mean of four observation in three wheat varieties at Day 1 Pre dawn and afternoon and Day 2 Pre dawn and afternoon and Day 1 is 5 days after imposition of water stress stress and 1 day after imposition of heat stress as measured by PAM 2500, FluorPen FP100 and MultispeQ. The quantum yield of primary photochemistry did not vary significantly between the instruments used in Control, Heat and Heat +Water stress treatments, however, in general the FluorPen FP100 showed higher absolute values in all the treatments, Heat +Water stress treatment exhibited significant difference in maximum quantum yield of primary photochemistry that Control, Water stress and Heat in all the instruments. Heat treatment did not show significant difference in the values of quantum yield as compared to control. The maximum recorded was 0.731 by Fluor Pen FP 100 in control as against 0.705 and 0.685 in PAM 2500 and MultispeQ respectively. Values recorded in Control, Heat and Heat +Water stress showed very little difference between instruments, on the other hand values recorded for water stress treatment showed a difference of 0.1 between Fluor Pen FP 100 (0.611) and MultispeQ and PAM 2500 with a mean of 0.519 and 0.510 respectively. Heat + Water stress recorded the lowest maximum quantum yield of primary photochemistry among all the treatments in all the instruments with 0.341, 0.353, 0.341 in MultispeQ, PAM 2500 and FluorPen FP100 respectively.

**Fig. 1.**
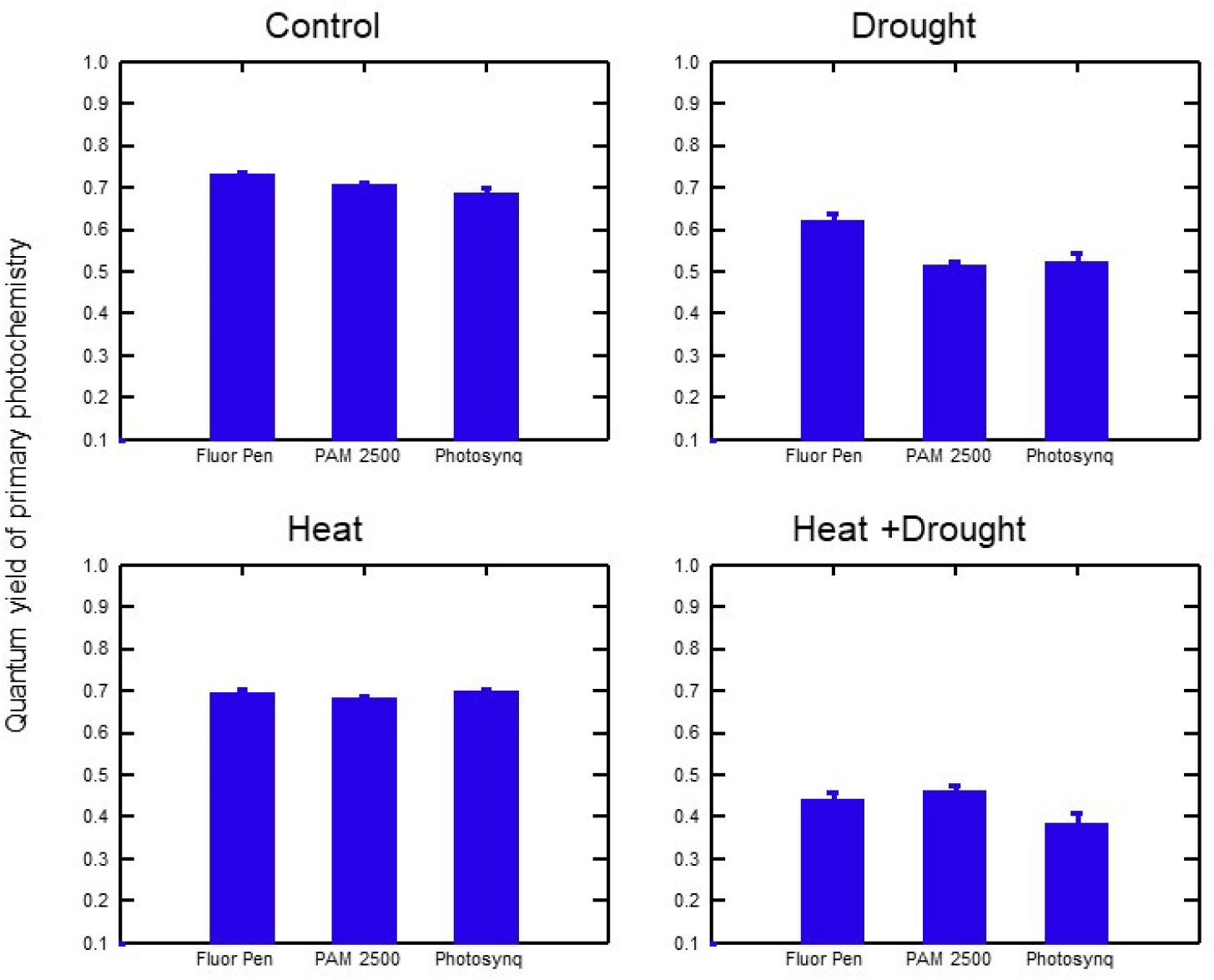
Maximum quantum yield of primary photochemistry in wheat under stress treatments measured by various instruments (vertical bars represent SEM). Values are mean of four observation in three wheat varieties at Day 1 Pre dawn and afternoon and Day 2 Pre dawn and afternoon and Day 1 is 5 days after imposition of drought stress and 1 day after imposition of heat stress

Maximum quantum yield of primary photochemistry in wheat under stress treatments measured by various instruments at different observation times in three wheat varieties is shown in Fig.2. The values are mean of three varieties Federation, Mace and Bolac. There were significant difference in the quantum yield at different times of observation with Day 2 afternoon showing the minimum values in all the instruments. Day 1 Predawn values showed significant difference between Heat + Water stress and the rest of the treatments in FluorCam FP 100 and MultispeQ, in the case of values recorded by PAM 2500 it was seen that both Water stress and Heat +Water stress treatments varied significantly from Control and Heat treatment. On the other hand in Day 1 afternoon the values recorded in Water stress and Heat +Water stress were significantly different from Control and heat treatments in all the instruments. A minimum value of 0.3 was recorded in both FluorCam FP 100 and MultispeQ in Heat +Water stress treatment. The values for Maximum quantum yield of primary photochemistry recorded by PAM 2500 did not vary significantly between Heat and Heat +Water stress treatments which was not the case in the other two instruments FluorCam FP 100 and MultispeQ. The observations taken on Day 2 Pre Dawn exhibited a uniform pattern in all the instruments with treatments Control and Heat showing significant difference from Water stress and Heat +Water stress in all the instruments. In Day 2 afternoon the treatments Water stress, Heat and Heat +Water stress recorded values of Maximum quantum yield of primary photochemistry which was significantly lower than that of control in all the instruments used with Heat +Water stress showing values of 0.4,0.2 and 0.2 in FluorPen FP 100, PAM 2500 and Multispeq respectively. All the instruments used showed a similar pattern in quantum yield of wheat plants under different treatments as was seen in Day 2 pre - dawn.

**Fig. 2.**
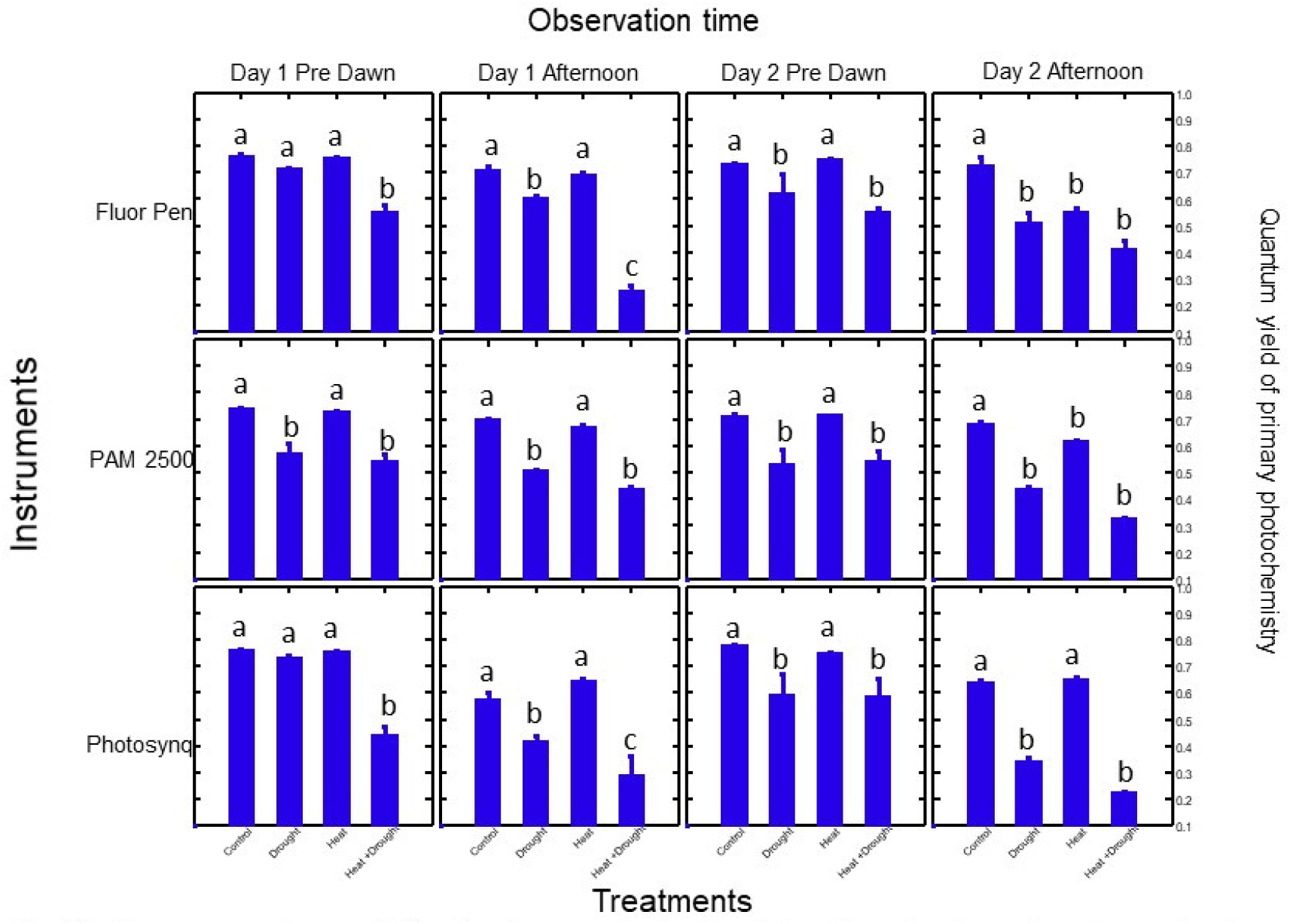
Maximum quantum yield of primary photochemistry in wheat under stress treatments measured by various instruments at different observation times. Day 1 is 5 days after imposition of drought stress and 1 day after imposition of heat stress (vertical bars represent SEM). Values of mean of three varieties of wheat, different letters indicate significance at 0.05

Fig.3 shows the Maximum quantum yield of primary photochemistry in wheat under stress treatments measured by various instruments in different varieties. Day 1 is 5 days after imposition of water stress stress and 1 day after imposition of heat stress. Values are mean of four observation in three wheat varieties at Day 1 Pre - dawn and afternoon and Day 2 Pre - dawn and afternoon. The values of quantum yield in all the varieties showed a similar pattern in all the instruments expect for Bolac measured by FluorPen. In the variety Bolac, PAM 2500 and Multispeq recorded values where the treatments Water stress and Heat +Water stress was significantly different from Control and Heat, this was not the case in FluorPen FP 100 where the treatment Water stress +Heat was significantly lower than that of the rest of the treatments viz., Control, Heat and Water stress. On the other hand all the instruments recorded a similar pattern of Maximum quantum yield of primary photochemistry in the varieties Federation and Mace. The difference in quantum yield between Control and Heat +Water stress was 0.3 recorded in PAM 2500 and FluorPen FP 100 respectively in variety Federation, on the other hand MultispeQ recorded a difference of 0.5, in the case of FluorPen FP 100 and MultispeQ in the variety Mace it was seen that there was a significant difference in the values of quantum yield between treatments Water stress and Heat +Water stress which was not the case in PAM 2500. Bolac showed higher values for Maximum quantum yield of primary photochemistry recorded by all the instruments in all the treatments as compared to the other varieties Federation and Bolac. The treatments Water stress and Heat +Water stress exhibited significantly lower values of quantum yield as compared to the treatments Control and Heat in all the varieties measured by all the instruments.

**Fig. 3.**
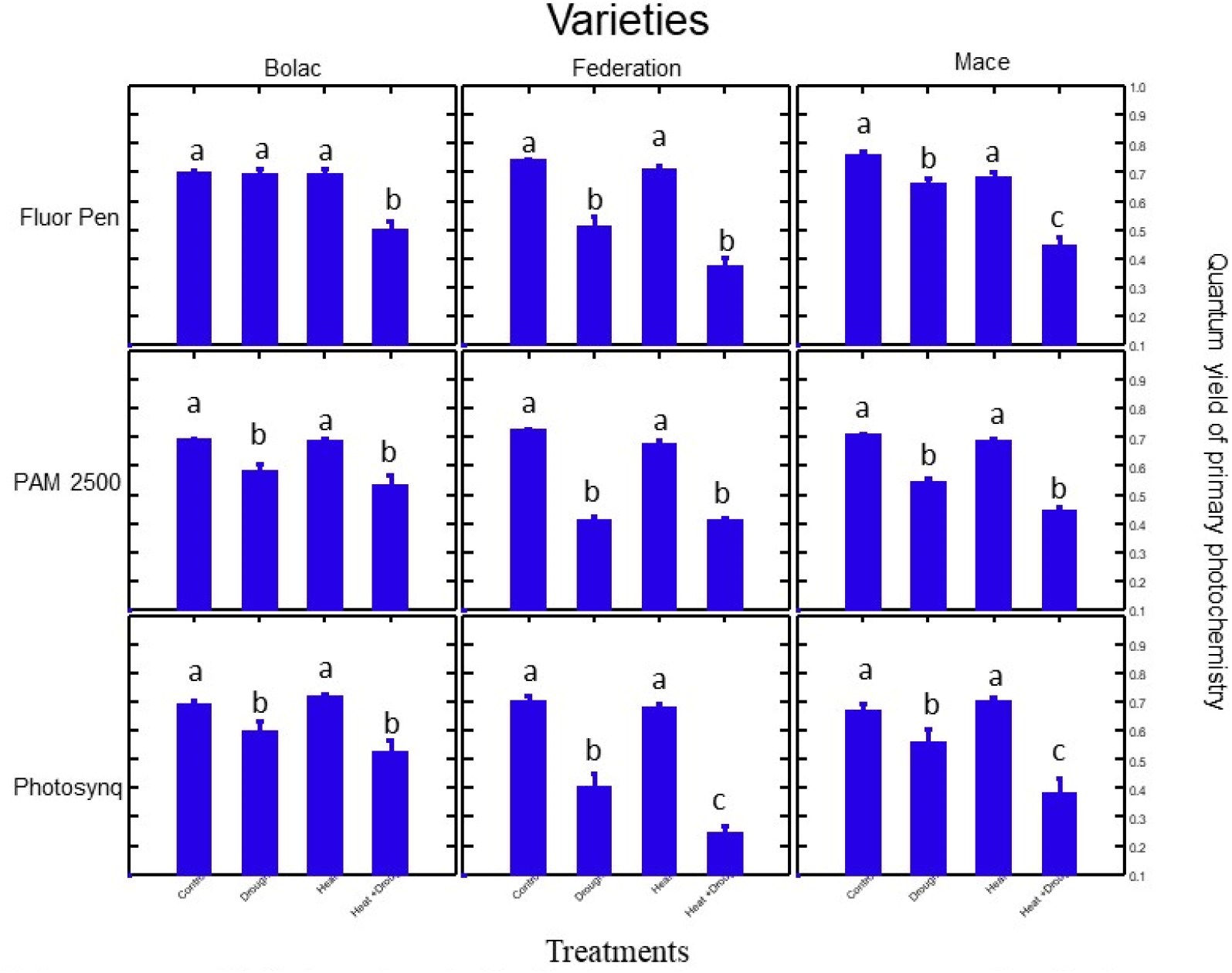
Maximum quantum yield of primary photochemistry in wheat under stress treatments measured by various instruments in different varieties. Day 1 is 5 days after imposition of drought stress and 1 day after imposition of heat stress (vertical bars represent SEM). Values are mean of four observation in three wheat varieties at Day 1 Pre dawn and afternoon and Day 2 Pre dawn and afternoon, different letters indicate significance at 0.05.

**Fig. 4.**
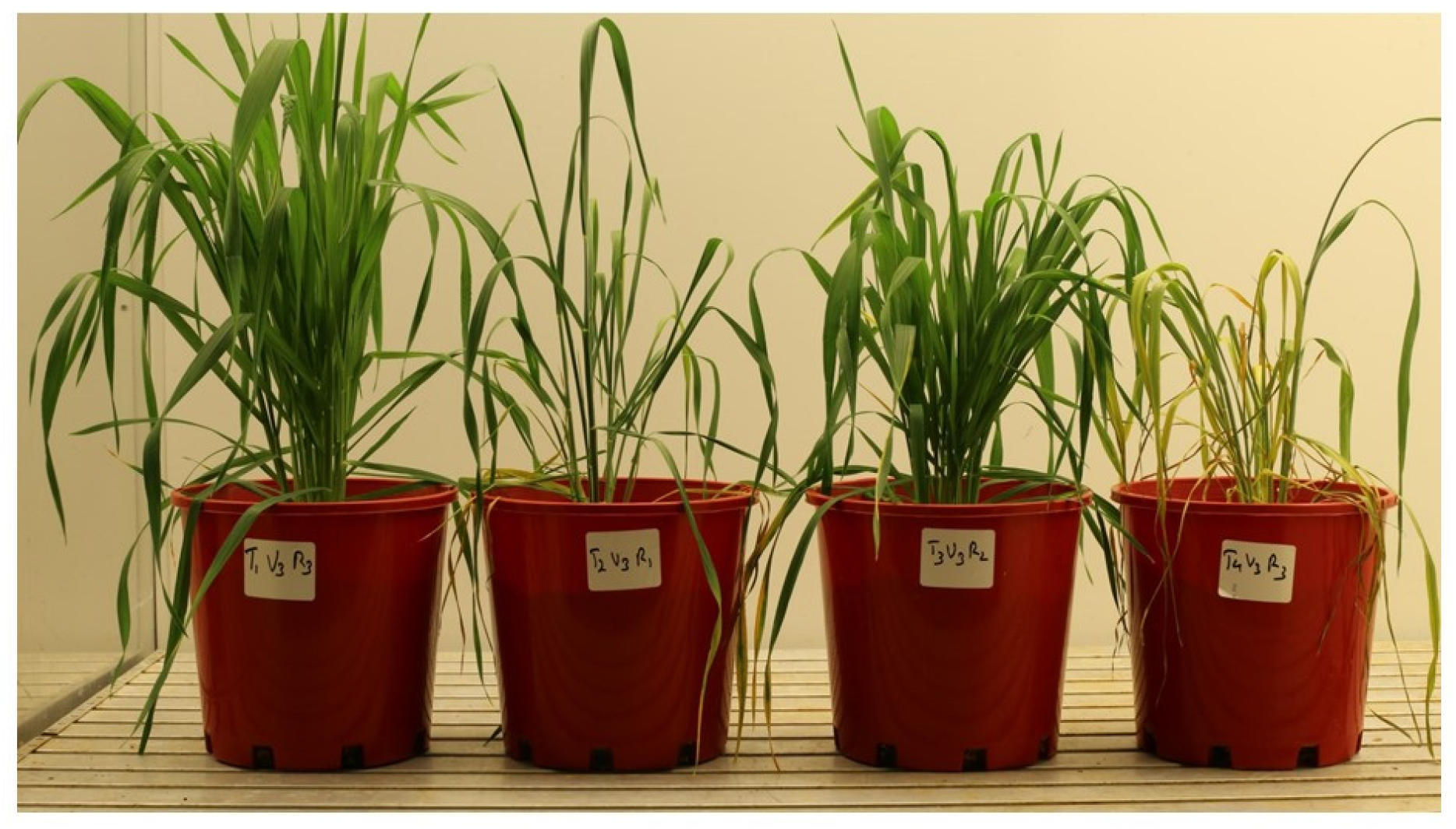
Wheat variety Bolac under different stresses - Control, Water Stress, Heat and Heat+ Water Stress

**Fig. 5.**
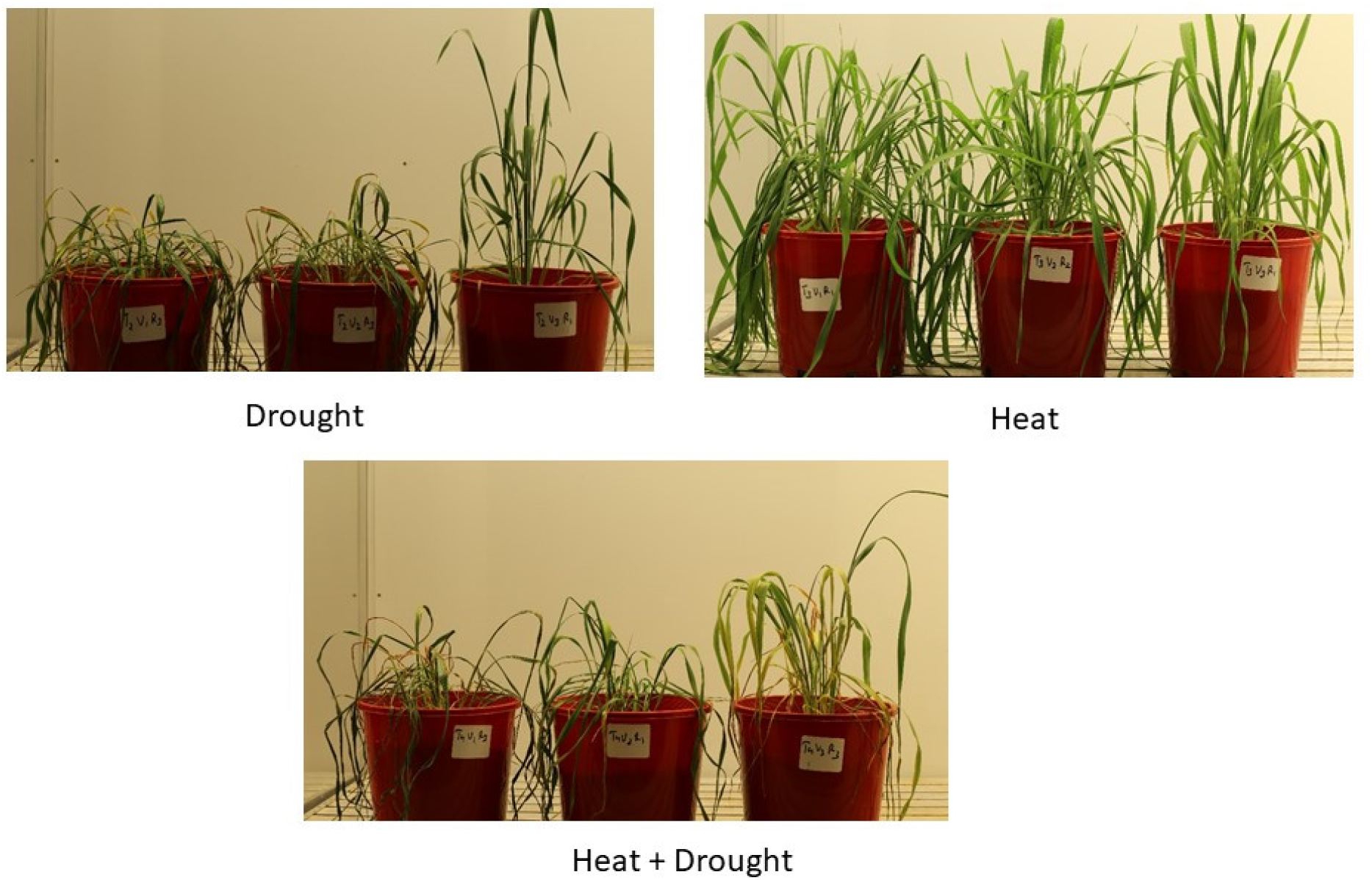
Differences in response to stress by wheat varieties-Federation, Mace, Bolac

**Fig. 6.**
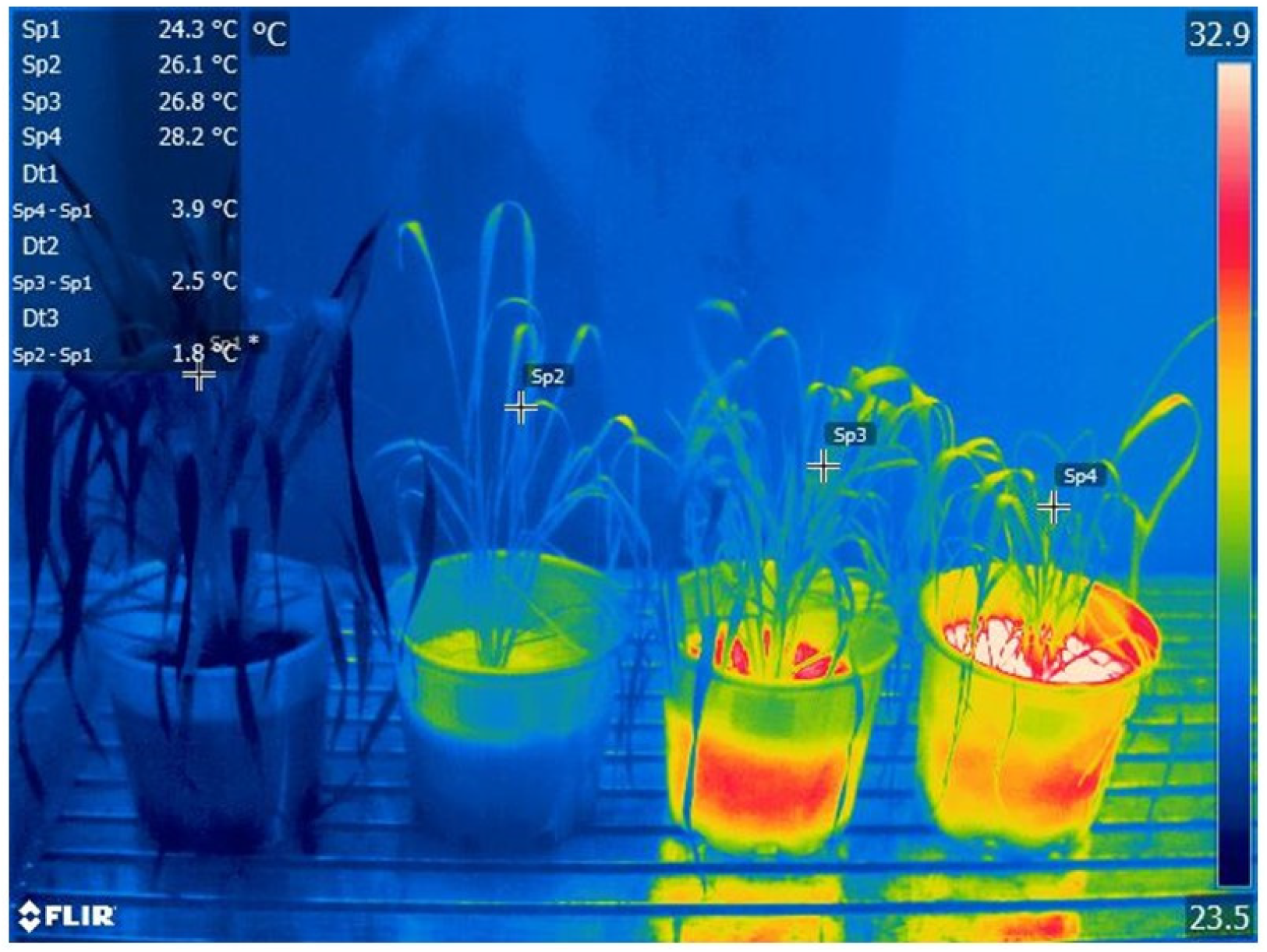
Infra Red Images of Wheat variety Bolac under different stresses - Control, Water Stress, Heat and Heat+Water Stress

**Fig. 7.**
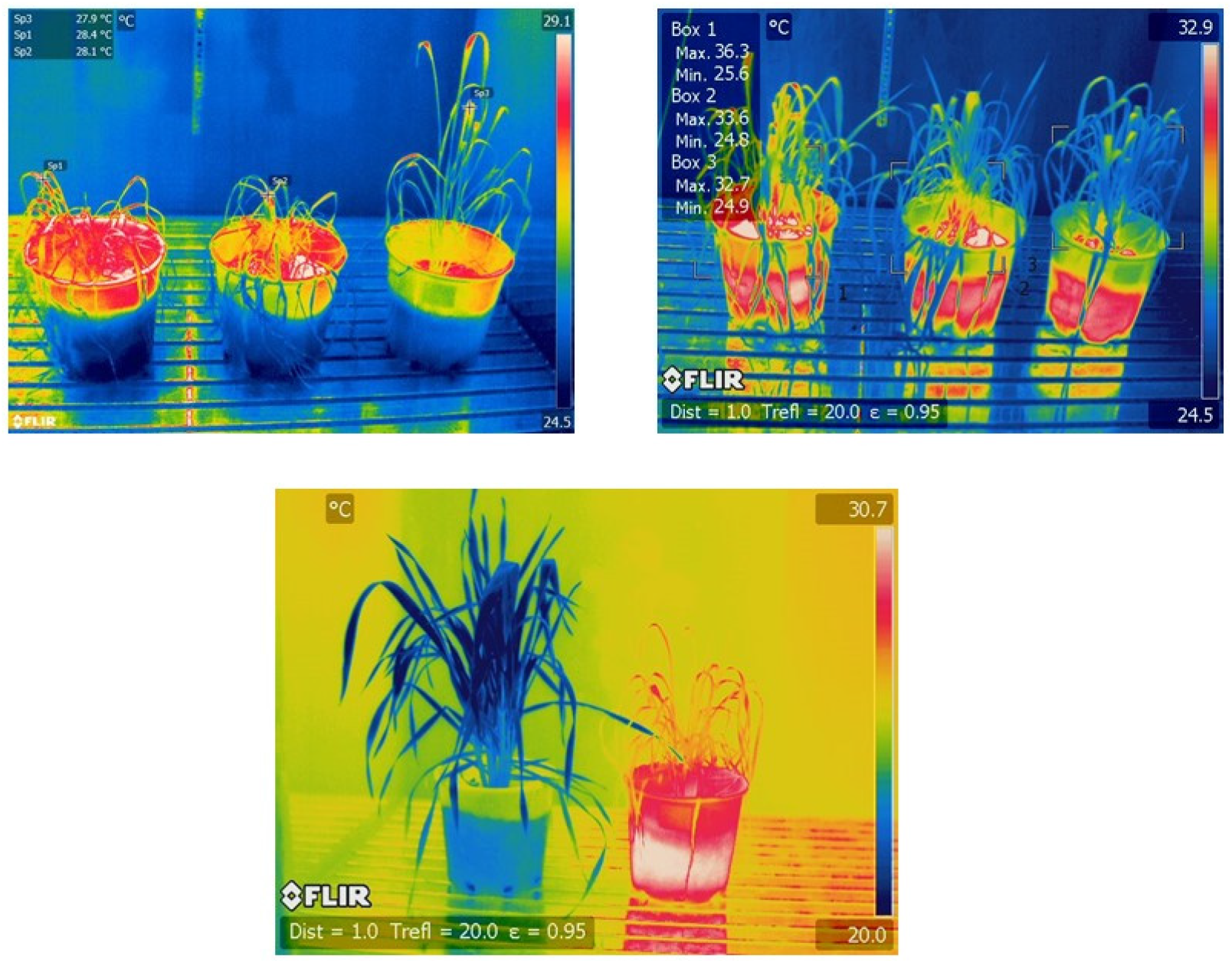
Infra Red Images of response to stress by wheat varieties - Federation, Mace, Bolac

## Discussion

Plants in general have a machinery of photosynthesis system that can experience the effects of many stresses at the same time and the response is more involving same mechanism by which protection and adaptation is achieved (Kreslavski et al 2013; Allakhverdiev et al 2008) A distinct K-step in the OJIP curve was seen in heat stressed plants which is attributed mainly to inhibition of the oxygen evolving complex in the PSII, this was not seen in the water deficit plants, which has been reported by Duarte et al (2015). K Step under heat stress is also reported by Rodriguez, et al (2015). We observed in our study that water deficit and heat stress both decreased the photosynthetic efficiency and capacity, driven by a loss in the connectivity between PSII antennae. Reaction Centre (RC) closure net rate and density increased, improving the energy fluxes entering the PS II, in spite of the high amounts of energy dissipated and the loss of PS II antennae connectivity. PI Performance index on absorption basis, incorporating the steps of antenna, reaction centre and electron transport parameters was lower than control in water deficit treatment followed by heat stress, similar results have been reported by Duarte et al (2017).

Heat +Water stress stress was more detrimental as compared to Heat or Water stress alone in terms of maximum quantum yield of photochemistry. Decrease in fluorescence at the J, I, and P level of the Kautsky curves point on either donor or acceptor site limitation at PS II. This could have been due to higher decrease in connectivity between PSII and its antennae resulting in lower photosynthetic efficiency. This resulted in the impairment and disruption of the electron transport. The Step before the OJIP observed in heat stress and heat +Water stress stress could be because of damage to Oxygen Evolving Complex indicating that low thermostability of the complex. The stress treatments had a reduction in the plastoquinone pool size as indicated by the reduced area above the OJIP curve. Our study indicated that PAM 2500 sensed both stresses separately and combined earlier than the other instruments, so in terms of sensitivity PAM 2500 was more effective than FluorPen FP 100 and MultispeQ. Rapid screening of stress was more effectively with FluorPen FP 100 and MultispeQ than by PAM 2500.

## Acknowledgement

AKS acknowledges the Department of Education and Training, Government of Australia for having given the Endeavour Executive Fellowship to undertake the study at High Resolution Plant Phenotyping Centre, CSRIO, Canberrra, AKS acknowledges CSIRO, Canberra, Australia for having supported my proposal for Endeavour Executive Fellowship. AKS acknowledges Indian Council of Agriculture Research, New Delhi, India and Director, Central Research Institute for Dryland Agriculture for permission to take up deputation to CSIRO for the study.

